# In lupus nephritis, specific *in situ* inflammatory states are associated with refractory disease and progression to renal failure

**DOI:** 10.1101/2021.09.03.458909

**Authors:** Rebecca Abraham, Madeleine Durkee, Junting Ai, Margaret Veselits, Gabriel Casella, Yuta Asano, Anthony Chang, Kichul Ko, Charles Oshinsky, Emily Peninger, Maryellen Giger, Marcus R. Clark

## Abstract

In human lupus nephritis (LN), tubulointerstitial inflammation (TII) on biopsy predicts progression to end stage renal disease (ESRD). However, while approximately half of patients with moderate or severe TII develop ESRD, half do not. Therefore, we hypothesized that TII is heterogeneous, with distinct inflammatory states associated with different renal outcomes. We interrogated renal biopsies from LN longitudinal and cross-sectional cohorts using both conventional and highly multiplexed confocal microscopy. To accurately segment cells across whole biopsies, and to understand their spatial relationships, we developed unique computational pipelines by training and implementing several deep learning models and other computer vision techniques. Surprisingly, across biopsies, high B cell densities were strongly associated with protection from ESRD. In contrast, elevated CD4-T cell population densities, which included CD8, γδ and double negative (CD4-CD8-δ-, DN) T cells, were associated with both acute refractory renal failure and gradual progression to ESRD. Interestingly, lymphocytes and dendritic cells were organized into discrete clusters or neighborhoods that could be characterized by the enrichment for specific cell populations. B cells were often organized into large neighborhoods with CD4+ T cells including T follicular helper-like cells. In contrast, the CD4-T cell populations formed small cellular neighborhoods whose frequency predicted subsequent progression to ESRD. These data reveal that in LN, specific *in situ* inflammatory states are associated with refractory disease and progression to ESRD.

**One sentence summary:** Using deep machine learning to analyze confocal microscopy data, we demonstrate that in lupus nephritis, CD4-T cell populations, including CD8+ and γδ T cells, organize into specific spatial neighborhoods that predict progression to renal failure.

## INTRODUCTION

For over 50 years, systemic lupus erythematosus (SLE) has been thought to result from a break in systemic tolerance and production of pathogenic autoreactive antibodies (*1, 2*). This canonical model is based on extensive studies of patient blood and spontaneous SLE-like animal models (*3, 4*). In the kidney, the manifestation of systemic autoimmunity is glomerulonephritis (GN). Indeed, LN is usually equated with GN (*5*). However, tubulointerstitial inflammation (TII)—and not GN—predicts progression to end stage renal disease (ESRD) (*6-9*).

Lupus TII is associated with a local immune response very different than the inflammation observed in glomeruli. Indeed, while TII is associated with infiltrating B cells, plasma cells, T follicular helper (Tfh) cells, plasmacytoid dendritic cells (pDCs) and myeloid dendritic cells (mDCs), these cells are rare in LN glomeruli (*10-15*). The immune cells found in TII are often organized into lymphoid-like architectures associated with local antigen-driven B cell clonal selection (*14, 15*). Therefore, in TII there are complex, intrinsic immune landscapes associated with progressive renal injury. There is a compelling need to understand *in situ* immunity in human lupus nephritis.

An initial roadmap to the lupus kidney was provided by the Accelerating Medicines Partnership (AMP)-funded single cell RNA-Sequencing (scRNA-Seq) of cells sorted from LN biopsies (*16, 17*). While these AMP investigations are informative, there are several limitations. The patient sample was small, and the frequency of each immune cell population has not been related to relevant histological features. A larger deficiency of scRNA-Seq is that all spatial information is lost. We do not know how populations spatially relate to each other in the kidney. This lack of spatial information prevents potential functional relationships from being identified.

Previously in lupus TII, we have used conventional immunofluorescence microscopy coupled to evolving computational and machine learning approaches to characterize the frequency of specific cell populations and identify cell:cell behaviors indicative of cognate immunity (*12, 13, 18*). However, a systemic analysis of TII and identification of prognostic features has been historically impeded by the complexities of analyzing immunofluorescence data from chronically inflamed kidneys, including tissue autofluorescence due to scarring, antibody cross-reactivity and patient heterogeneity. Artificial intelligence algorithms have led to significant progress in automated detection and analysis of cells in confocal images (*19-22*). These approaches have been used successfully in cancer biopsy image analysis (*23*). However, conventional methods for cell detection and segmentation are not easily generalized to chronically inflamed organs.

Herein, we describe multiple computational pipelines employing deep learning algorithms that provide high-throughput assessments of cell phenotypes and cellular architectures. These methods have been developed and validated in LN image datasets consisting of both discrete fields of view and entire biopsy sections images. Integration of these data reveal that CD4-T cell populations, comprised of CD8+, γδ and double negative (CD4-CD8-δ-, DN) T cells, often organized into small cellular neighborhoods, are both associated with acute refractory disease and predict progression to renal failure. In contrast, regions of high B cell density were associated with patients who did not progress to renal failure. These and other findings indicate that systemic and *in situ* autoimmune pathogenic mechanisms are different in LN, and each might require specific targeted therapies.

## RESULTS

### Accurate segmentation of immune cells in lupus nephritis kidney biopsies

To probe the relationship between TII and clinical outcome we used a well-characterized cohort of 55 biopsy-proven lupus nephritis patients with at least two years of follow up (**Supplementary Table 1**). Within this cohort, 19 patients progressed to end stage renal disease (ESRD+) while 36 did not (ESRD-). The ESRD+ and ESRD-groups did not differ in length of follow-up, duration of disease, or patient age (**Supplementary Fig. 1A-C**). Additional information about patient treatment can be found in Supplementary Table 1. Thirty-eight patients had moderate or severe TII distributed across both outcome groups. Based on previous studies (*6*), we hypothesized that differences in renal outcome would be related to differences in *in situ* adaptive immunity, such as frequency and organization of principal cellular effectors. Therefore, we stained each biopsy for six markers, CD3, CD4, CD20, CD11c, BDCA2, and DAPI to characterize five classes of immune cells: CD3+CD4+ T cells, CD3+CD4-T cells, CD20+ B cells, BDCA2+ plasmacytoid dendritic cells (pDCs), and CD11c+ myeloid dendritic cells (mDCs). Across the 55 biopsies, we captured all regions of interest (ROIs) with detectable CD3+ T cells, resulting in 865 ROIs. Image ROIs were 1024 × 1024 pixels, with a pixel size of 0.1058 μm. These data are referred to as the high resolution (HR) dataset.

Lupus nephritis is often characterized by chronic and intense inflammation in which accurate cell segmentation can be difficult due to the high cell densities and structured background signal (*12, 21*). Therefore, we trained deep convolutional neural networks (DCNNs) to perform automatic cell detection, classification, and segmentation (collectively known as instance segmentation) on the HR dataset. To achieve optimal performance across all cell classes, we split the five-class cell detection into two tasks: instance segmentation of lymphocytes and instance segmentation of DCs (**Fig. 1A**). For each task, a separate instance of a region-based DCNN architecture, Mask R-CNN, was independently trained (**Fig. 1B**) (*24*). Each Mask R-CNN was trained on 246 manually segmented images with a validation set of 65 manually segmented images used for hyperparameter tuning. On a test set of 30 images from patients unique to the training and validation data, the lymphocyte detection network had an F1-score of 0.75 and the DC detection network of 0.57 while the overall F1-score for detection of all 5 cell classes was 0.74, yielding excellent concordance (**Fig. 1C**). By implementing DCNNs, we achieve rapid and accurate multi-class instance segmentation.

**Fig. 1.**
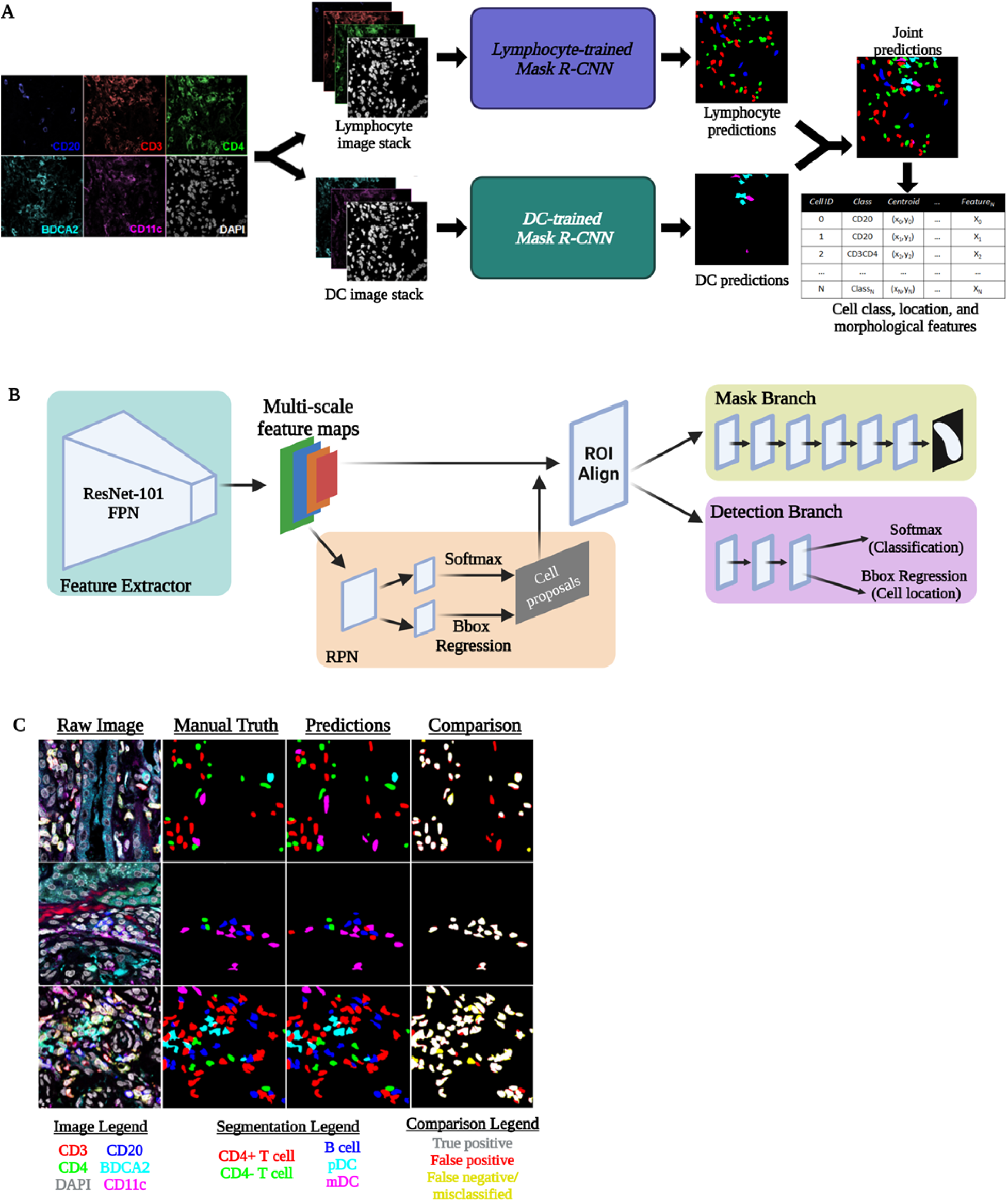
Instance segmentation of immune cells in high-resolution fluorescence microscopy images of LN kidney biopsies. A) Automatic instance segmentation of five immune cell classes was performed by combining predictions from two instances of Mask R-CNN: one trained to segment CD20+, CD3+CD4-, and CD3+CD4+ lymphocytes and one trained to segment pDCs and mDCs. Cell location, class, and morphological features were calculated from joint predictions. B) The Mask R-CNN architecture is comprised of a ResNet Feature Pyramid Network (FPN) backbone used for feature extraction, a region proposal network (RPN) used to generate cell proposals, and two parallel branches used for 1) semantic segmentation (mask branch), and 2) classification (softmax layer) and localization (bounding box (Bbox) regression) of cell proposals. C) Representative segmentations produced by the multi-network pipeline showed strong agreement with the expert-defined manual segmentations. Created with BioRender.com.

### Specific *in situ* immune cell densities associated with progression to renal failure

Automatic cell segmentations were used to describe and quantify the spatial distribution all five cell classes in the HR dataset. Comparison of overall cell densities (total cells/ROI) in ESRD-patients and ESRD+ patients revealed no significant differences (**Fig. 2A**). However, the total cell count per sample was higher in the ESRD+ cohort, reflecting larger overall areas of inflammation (**Fig. 2B**). In contrast to overall cell density, there were differences in the cellular constituents of inflammation between the two patient cohorts. Surprisingly, ROIs from ESRD-patients had higher densities of B cells relative to ROIs from ESRD+ patients (**Fig. 2C**). In contrast, ROIs from ESRD+ patients had increased densities of CD4-T cells (**Fig. 2D**). There were no significant differences in the densities of CD4+ T cells, pDCs, or mDCs between patient cohorts (**Fig. 2E-G**).

**Fig. 2.**
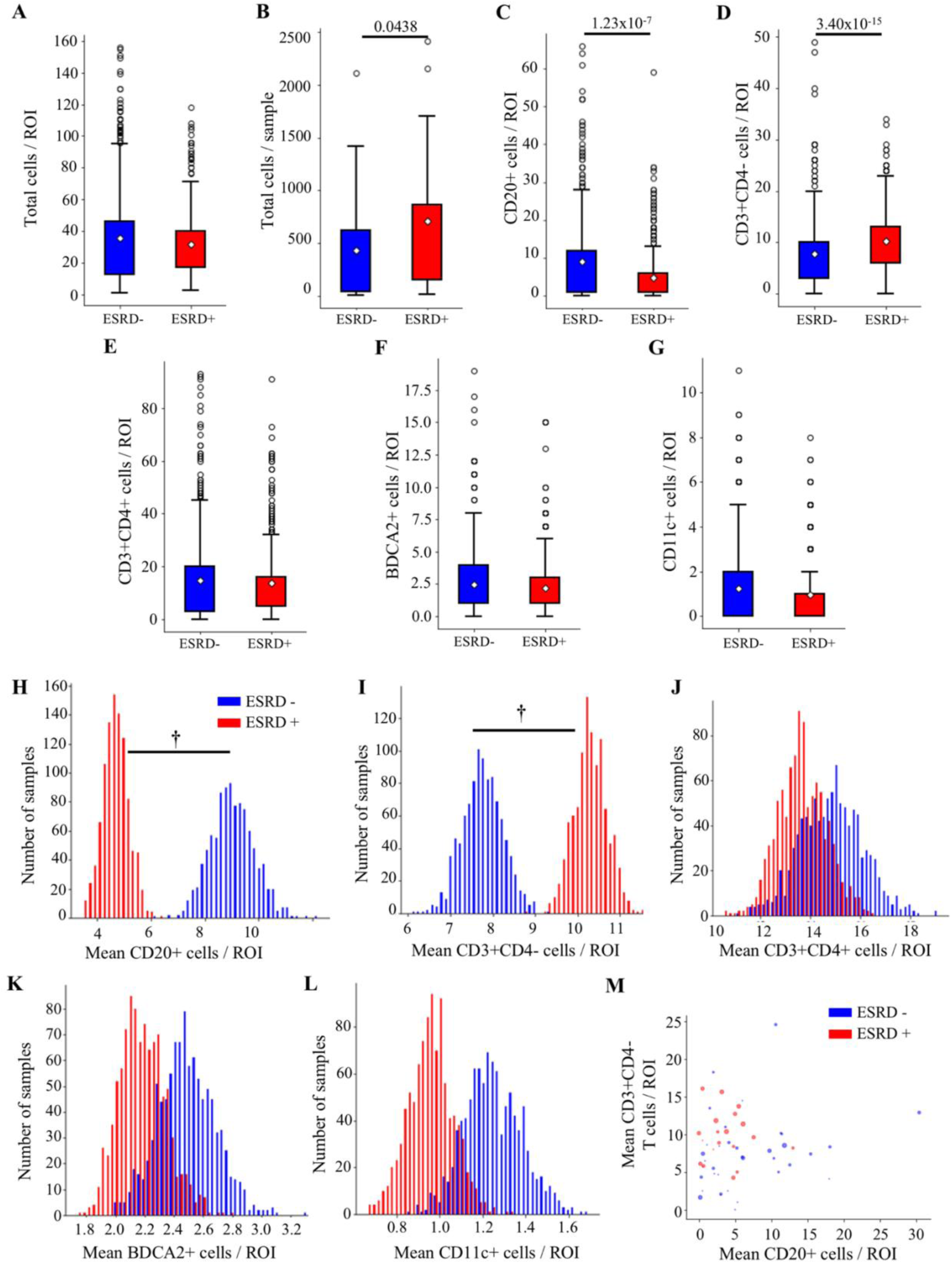
Higher CD4-T cell density and lower B cell density associated with progression to ESRD. A) Local cell density is compared for ESRD-patients (n=437 ROIs) and ESRD+ patients (n=428) for all cells. B) Total cells per patient grouped by ESRD status. Local cell density by cell class is compared between ESRD- and ESRD+ patient for C) CD20+ cells, D) CD3+CD4-cells, E) CD3+CD4+ cells, F) BDCA2+ cells, and G) CD11c+ cells. All cell density comparisons were done with a Mann-Whitney U Test with a Bonferroni correction for multiple comparisons, with significant p values denoted in each panel. Bootstrapped sample means of ESRD- (blue) and ESRD+ (red), ROIs for H) CD20+ cells/ROI, I) CD3+CD4-cells/ROI, J) CD3+CD4+ cells/ROI, K) BDCA2+ cells/ROI, L) CD11c+ cells/ROI. Average B cell and CD4-T cell count per ROI for each patient biopsy is shown in (M). Point size is weighted by the TI chronicity score for each patient. (**†** 95% confidence interval does not overlap with 0)

Although there were fewer ESRD+ patients, on average these patients had more ROIs captured per biopsy. To mitigate any effect from this class imbalance, we performed a bootstrapping analysis. The pools of ESRD+ and ESRD-ROIs were iteratively sampled with replacement 1000 times to produce samples of 200 ROIs from each group (ESRD+ and ESRD-). The distribution of mean cell densities between ESRD+ and ESRD-patients revealed distinct, non-overlapping peaks for both B cells and CD4-T cells (**Fig. 2H-I**). In contrast, there was substantial overlap in the distribution of sample means between ESRD+ and ESRD-patients for CD4+ T cells, pDCs and mDCs (**Fig. 2J-L**). The 95% confidence intervals of the difference in means between ESRD+ and ESRD-patients revealed for both B cells and CD4-T cells did not cross zero (**Supplementary Fig. 2A-B**). In contrast, the 95% confidence interval for the difference in means for the remaining cell types did cross zero (**Supplementary Fig. 2C-E**). These data indicate that the observed differences in B cell and CD4-T cell densities between ESRD+ and ESRD-patients are robust. Therefore, we conclude that high B cell densities are associated with a good prognosis while high densities of CD4-T cells are associated with progression to renal failure.

When we examine these densities on the patient level, we observe that in patients with high CD4-T cell densities, B cell densities tend to be low (**Fig. 2M**). As indicated by point size, these tended to be ESRD+ patients with higher TI chronicity scores. The converse appeared true, as patients with higher B cell densities tended to have low TI chronicity scores and be ESRD-. These data suggest that lupus TII is associated with two or more distinct inflammatory states, each associated with a different prognosis.

### Patients who present in renal failure have a skewed *in situ* inflammatory state

Within the ESRD+ group of patients was a small yet distinct cohort of five patients that either were in renal failure at the time of biopsy or progressed to renal failure within two weeks of biopsy collection. If these patients are treated as their own unique outcome group (ESRD current), differences in the density of specific cell classes become even more apparent (**Fig. 3**). There are progressively fewer B cells/ROI between ESRD-, ESRD+ and ESRD current groups, respectively (**Fig. 3A**). The opposite trend is observed for CD4-T cell densities (**Fig. 3B**). In contrast, there were no apparent differences in CD4+ T cells or pDCs in the ESRD current patients (**Fig. 3C-D**). Remarkably, there was a profound depletion of mDCs in the ESRD current cohort (**Fig. 3E**).

**Fig. 3.**
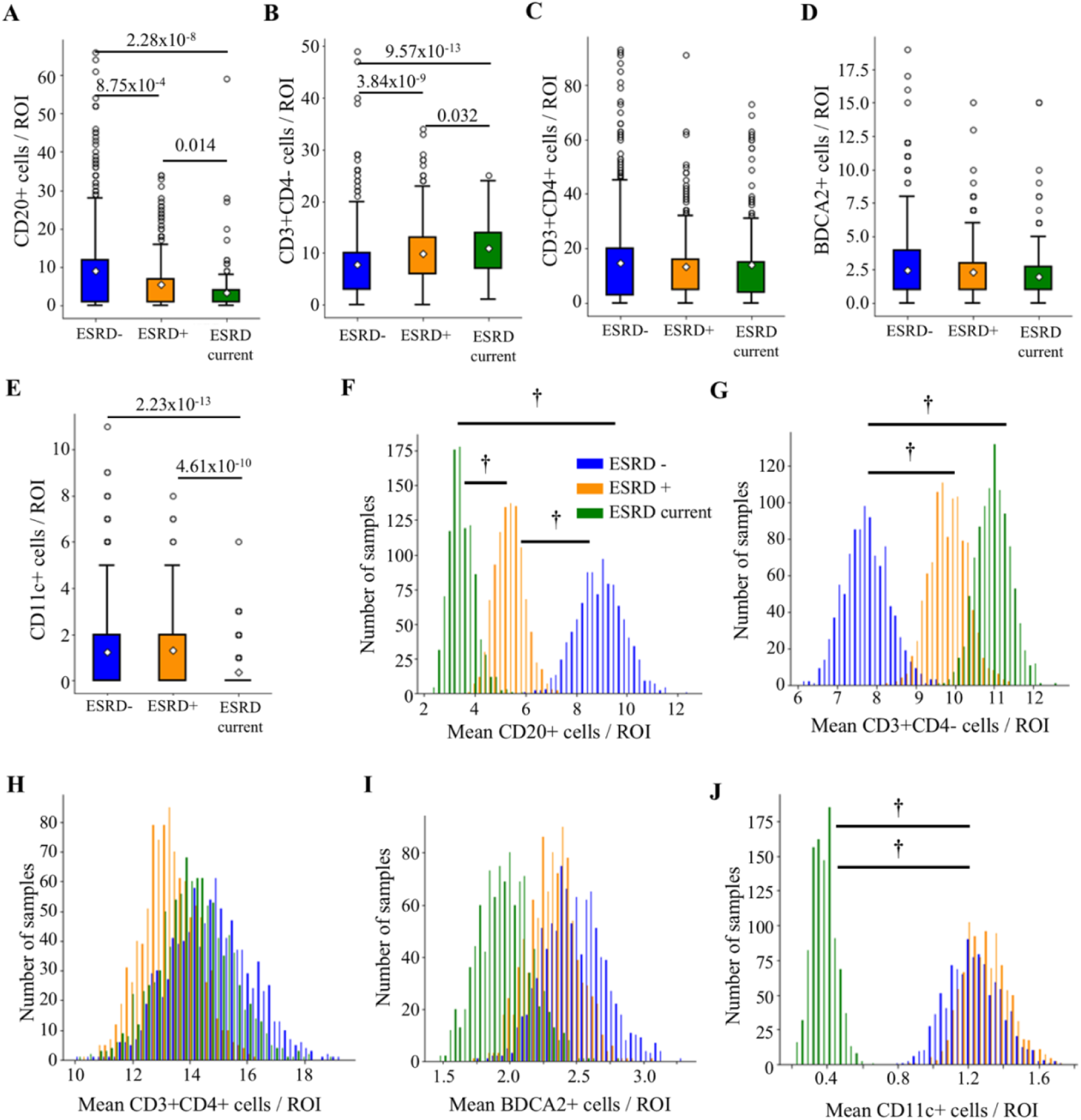
Local cell densities are associated with progressively worse renal outcomes. Local cell density is compared across ESRD-patients (n=437 ROIs), ESRD+ patients (n=266), and ESRD current patients (n=162) for A) CD20+ cells, B) CD3+CD4-cells, C) CD3+CD4+ cells, D) BDCA2+ cells, and E) CD11c+ cells. All cell density comparisons were done with a Mann-Whitney U Test with a Bonferroni correction for multiple comparisons, with significant p values denoted in each panel. Bootstrapped sample means of ESRD-(blue), ESRD+ (orange), and ESRD current (green) ROIs for F) CD20+ cells/ROI, G) CD3+CD4-cells/ROI, H) CD3+CD4+ cells/ROI, I) BDCA2+ cells/ROI, J) CD11c+ cells/ROI. (**†** 95% confidence interval does not overlap with 0)

A three-group bootstrapping analysis was performed to assess the effect of class imbalance in patient numbers. ESRD current patients had the lowest mean density of B cells, followed by ESRD+, with ESRD-having the highest density of B cells (**Fig. 3F**). Confidence intervals for the pairwise differences between bootstrapped samples did not overlap with zero (**Supplementary Fig. 3A**). An inverse, stepwise relationship was observed for CD4-T cells with progressively higher densities found in the ESRD+ and ESRD current relative to ESRD-cohorts (**Fig. 3G, Supplementary Fig. 3B**). ESRD-current patients were also well separated from the other two cohorts with respect to local mDC abundance (**Fig. 3J, Supplementary Figure 3E**). As expected, there were no differences between the three groups with respect to CD4+ T cells or pDCs (**Fig. 3H-I, Supplementary Fig. 3C-D)**. These findings indicate that LN patients that present in renal failure have a skewed inflammatory state with abundant CD4-T cells, relatively few B cells and a depletion of mDCs.

### Specific cellular neighborhoods associated with progressive and refractory renal disease

We next explored the relative *in situ* spatial relationships between the different immune cell classes. First, for every cell in the dataset, we identified the nearest neighbor using centroid to centroid distances. All cell classes except mDCs were significantly more likely to have a B cell as their nearest neighbor in ESRD-biopsies (**Fig. 4A**). In contrast, all cell classes were significantly more likely to have a CD4-T cell nearest neighbor in ESRD+ biopsies (**Fig. 4B**). Additionally, both B cells and CD4-T cells showed a strong propensity for co-localization with cells of the same type.

**Fig. 4.**
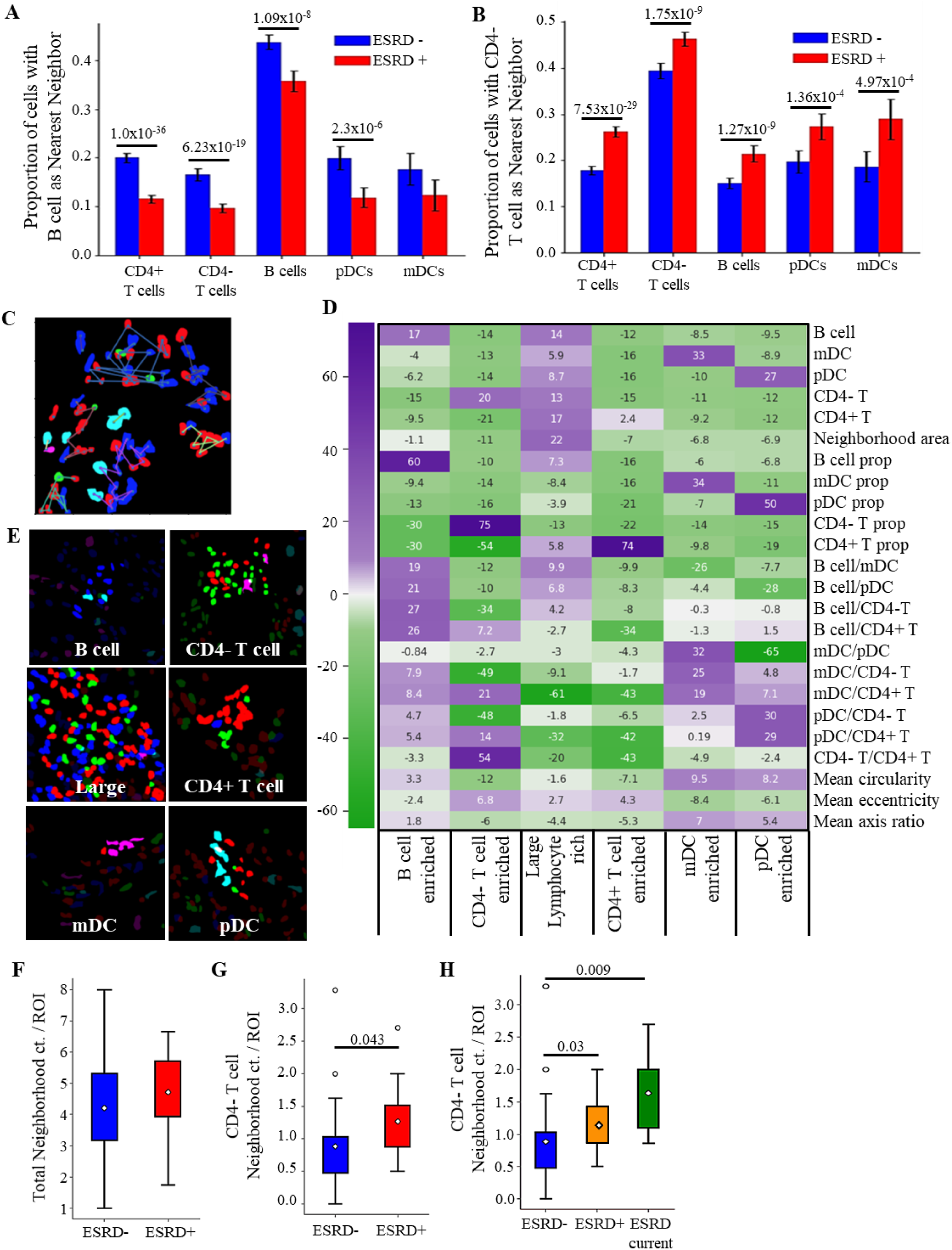
Specific cellular neighborhoods associated with renal failure. Proportions of cells that have A) CD3+CD4-T cells and B) CD20+ B cells as nearest neighbors in ESRD+ and ESRD-patients (chi-squared test for independence with Bonferroni correction for multiple comparisons). C) Representative neighborhoods detected by DBSCAN. D) Heatmap showing test statistics for each feature from leave-one-out t-tests used to define six types of cell neighborhoods, colored by the magnitude of the test statistic. E) Representative neighborhoods from each defined class. The abundance of neighborhoods between the patient cohorts, normalized by the number of ROIs per patient, is compared by Mann-Whitney U Test with a Bonferroni correction for F) all cell neighborhoods and G) CD4-T cell neighborhoods. A 3-group comparison for CD4-neighborhoods, splitting the ESRD+ population into ESRD+ and ESRD current patients is shown in (H). Significant p-values after correcting for multiple comparisons are denoted in each panel.

Local cellular organization was then probed by grouping cells into spatially discrete neighborhoods. DBSCAN, a density-based clustering algorithm (*25*), was implemented to define cell neighborhoods using a maximum intercellular centroid-centroid distance. Variation in this maximum distance between 50 and 150 pixels resulted in a range of neighborhood sizes varying between those that contained just a few cells (50 pixels) to those that encompassed large areas of inflammation (150 pixels) (**Fig. 4C, Supplementary Fig. 4A**). A maximum distance of 100 pixels (∼10.6 µm) was selected, as this distance approximates a cell body and appeared to capture observable groupings of cells across the dataset.

Using this 100-pixel cutoff and a minimum neighborhood size of two, DBSCAN detected 4022 cell neighborhoods. Each neighborhood was characterized by a set of 24 quantitative features including cell type frequency, cell type proportion, ratios of cell types, cell shape features, and neighborhood area (**Supplementary Fig. 4B**). K-means clustering was then applied to define classes of neighborhoods, with k=6 classes determined ideal by bootstrapping cluster descriptors including the within cluster sum of squares (WCSS) and the delta WCSS (**Supplementary Fig. 4C-D**). The test score from a leave-one-out t-test approach was used to determine which features or combination of features best distinguished the six neighborhood groups (**Fig. 4D**). The most distinctive feature(s) for each group was used to describe the cell neighborhoods as follows: 1) B cell enriched cluster, 2) CD4-T cell enriched cluster, 3) Large, lymphocyte enriched cluster, 4) CD4+ T cell enriched cluster, 5) mDC enriched cluster, and 6) pDC enriched cluster (**Fig. 4E**).

Tertiary lymphoid structures (TLSs) have been previously identified in the context of lupus nephritis (*15*). Although we cannot explicitly define TLSs in this dataset, we hypothesized that some of the large, lymphocyte enriched neighborhoods might approximate TLSs. For example, we noted that within this group, 28.6% of the cells were B cells and 48.3% were CD4+ T cells. 96.1% of these neighborhoods met the following criteria: 1) contained at least 20 cells, 2) both B cells and CD4+ T cells were represented in the neighborhood and 3) at least 50% of all cells were B cells and/or CD4+ T cells. Therefore, the vast majority of large, lymphocyte enriched neighborhoods, have features consistent with TLSs.

We then examined how these six classes of neighborhoods were distributed between the ESRD-and ESRD+ patients. After normalizing by the number of ROIs captured for each patient, ESRD- and ESRD+ patients had no difference in their total neighborhood count per ROI (**Fig. 4F**). However, ESRD+ patients had significantly higher prevalence of CD4-T cell enriched neighborhoods relative to the ESRD-patients (**Fig. 4G**). The per ROI prevalence of the other classes of neighborhoods did not correlate with renal outcome (**Supplementary Fig. 5A-E)**. We next examined if the CD4-neighborhoods differed between the ESRD-, ESRD+ and ESRD current patient groups (**Fig. 4H**). ESRD+ and ESRD current patients had a statistically higher prevalence of neighborhoods from the CD4-cluster than ESRD-patients These data demonstrate that, on a per patient basis, the prevalence of small, CD4-T cell enriched neighborhoods is strongly associated with progressive renal disease.

### Cell detection and segmentation in highly multiplexed, full-biopsy images

To better characterize *in situ* lymphocyte populations, we performed highly multiplexed confocal microscopy on a separate dataset of 18 lupus nephritis biopsies. In this highly multiplexed (HMP) dataset, we interrogated a set of nine markers (CD3, CD4, CD8, ICOS, PD1, FoxP3, CD20, CD138, and DAPI). This highly multiplexed panel was obtained using four-color confocal microscopy and iterative stripping and reprobing (*26*). Additionally, we imaged full biopsy sections rather than capturing isolated ROIs, thereby facilitating a more complete and unbiased spatial analysis.

Full biopsy images for all stains were aligned with the DAPI channel (**Fig. 5A**). Two new instances of Mask R-CNN were trained to perform single-marker and dual-marker instance segmentation (**Fig. 5B**). Briefly, ROIs from the HR dataset (pixel size = 0.1058 μm) were broken up into 512 × 512-pixel tiles to pre-train each Mask R-CNN. Each network was then fine-tuned using small sets of manually segmented 512 × 512-pixel tiles from the HMP dataset (pixel size = 0.221 μm). The single-marker Mask R-CNN was used to predict B cells (CD20+) and plasma cells (CD138+), while the dual-marker Mask R-CNN was used to predict single positive and double positive T cells. The three main classes of T cells were determined by combining predictions from a CD3/CD4/DAPI image stack with predictions from a CD3/CD8/DAPI image stack at the same location in the tissue: CD4+, CD8+, and CD4-CD8- (DN) (**Fig. 5C**). The dual-marker Mask R-CNN was also used to generate cell predictions on CD3/ICOS/DAPI and CD3/PD1/DAPI images. The resulting single-positive (CD3+ICOS- or CD3+PD1-) and double-positive (CD3+ICOS+ or CD3+PD1+) predictions were to define ICOS and PD1 expression for every putative T cell in the dataset. FoxP3 images were binarized by thresholding individual image tiles. T cell predictions with > 25% overlap with this binary mask were determined to be FoxP3+.

**Fig. 5.**
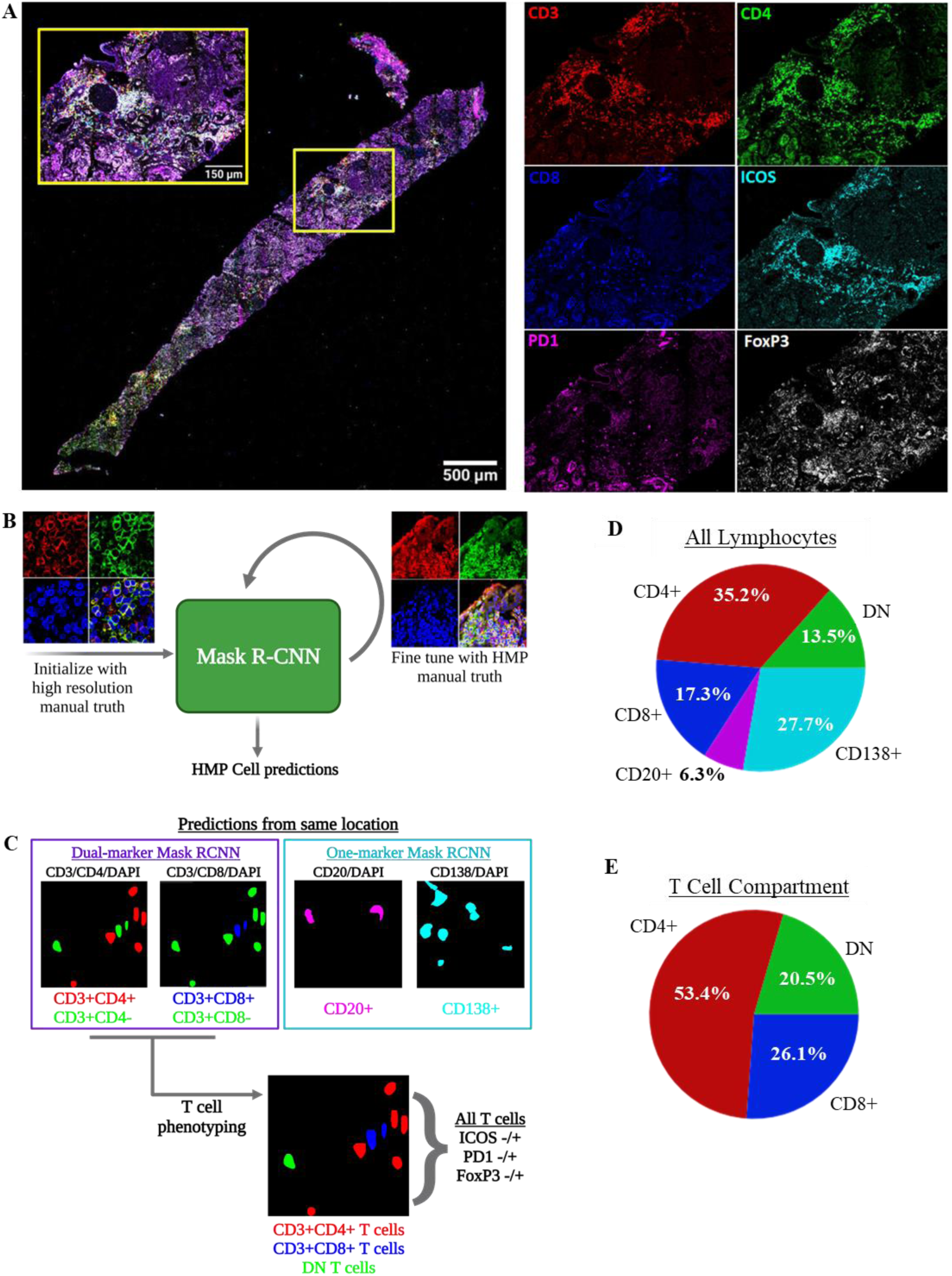
Cell detection, segmentation, and phenotyping in highly multiplexed fluorescence microscopy images. A) Representative composite of a full biopsy section, shown with merged and with isolated panels of CD3, CD4, CD8, ICOS, PD1, and FoxP3. B) Schematic of procedure for training and fine-tuning a Mask R-CNN for instance segmentation of cells in highly multiplexed microscopy images. C) Dual-marker and single-marker cell predictions are used to establish base lymphocyte classes. All T cell predictions are further described by ICOS, PD1, and FoxP3 expression. D) Breakdown of frequencies of the five base classes in the HMP dataset. E) Frequencies of CD4, DN and CD8 T cells within the T cell compartment. Figure 5A-C created with BioRender.com.

### CD4-T cells contain CD8, γδ and other DN T cell populations

T cells comprised over 65% of predicted lymphocytes in the HMP dataset (**Fig. 5D**). Plasma cells were the second most abundant class, comprising ∼28% of detected lymphocytes. B cells were least prevalent at only ∼6%. CD4+ T cells were the most abundant cell class across all five main classes, making up 35% of detected lymphocytes, and over 50% of detected T cells (**Fig. 5E**). Surprisingly, CD8+ T cells were only slightly more abundant than DN T cells, comprising ∼17% of detected lymphocytes and ∼26% of detected T cells.

To further characterize these DN T cells, we interrogated public scRNA-Seq data of immune cells infiltrating the kidney of lupus nephritis patients (*16*). We identified naïve T and CTL clusters in intrarenal immune cells by unsupervised clustering and canonical marker expression (**Fig. 6A**). Within these T cell clusters, 21% were DN, as measured by the UMI of *CD4, CD8A*, and *CD8B* (**Supplementary Fig. 6A**). Several T cell subtypes do not express CD4 nor CD8 including natural killer (NK) T cells and γδ T cells. Indeed, there was a small increase in *CD3D* in cells assigned to the NK cell class, suggesting a NKT phenotype. However, there was not a substantial enrichment for NKT cell markers in the DN subset (**Supplementary Fig. 6B**).

**Fig. 6.**
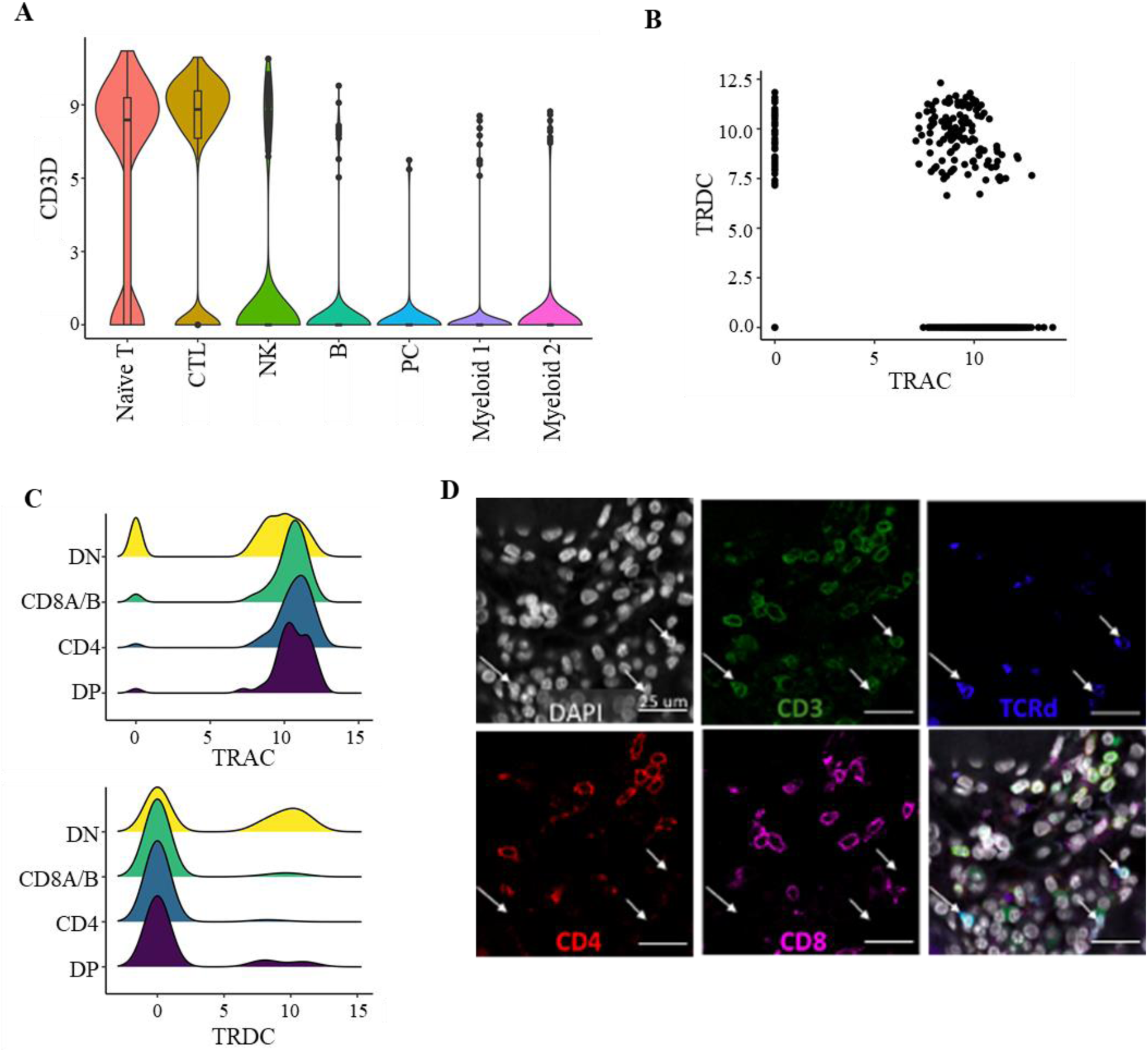
Identifying γδ T cells in LN. A) distribution of *CD3D* in cell clusters identified in scRNA-Sequencing data from LN kidney samples. B) Expression of *TRAC* and *TRDC* in T cells identified in scRNA-Seq data. C) Comparison of *TRAC* and *TRDC* expression in identified DN, CD8+, CD4+ and DP T cells. D) Representative image of double negative (CD4-CD8-)γδ (TCRd+) T cells in LN biopsy, marked by white arrows.

Next, we compared TCRα and δ chain expression (*TRAC* and *TRDC*). Some cells were apparently positive for both *TRAC* and *TRDC*, likely due to sequence homology between these genes (**Fig. 6B**). However, *TRAC*-cells and *TRDC*+ cells were both enriched in DN population (**Fig. 6C**). These results suggest that a portion of the DN T cells observed in lupus nephritis are γδ T cells. To further examine this possibility, we stained eight lupus nephritis biopsies with antibodies specific for CD3, CD4, CD8, TCRδ, and imaged 281 ROIs (**Fig. 6D**). Per biopsy, 51.4% ± 21.3% of DN T cells were positive for TCRδ. These findings indicate a substantial fraction of T cells in lupus nephritis do not detectably express CD4 or CD8, and approximately half of these DN T cells are γδ T cells.

### *In situ* distributions of exhausted, regulatory and helper T cell populations

We then examined the distributions of ICOS, PD1 and FoxP3 in the T cell subsets. Roughly 30% of CD8+ T cells in the HMP dataset are PD1 positive (**Supplementary Fig. 7A**) suggesting an exhausted phenotype. Approximately 25% of CD8+ T cells were “exhausted” by the definition of PD1+ICOS-FoxP3- (*27*). This is coherent with observations from murine lupus models in which exhausted tissue-infiltrating CD8+ T cells are relatively common (*28*). However, PD1 is only one marker of exhaustion and human lupus renal scRNA-Seq data suggests CD8+ T cell exhaustion is infrequent (*16*).

A surprisingly small percentage (5.41%) of CD4+ T cells were FoxP3 positive, while fewer still were also PD1-ICOS-, suggesting that Tregs comprise only about 2.5% of CD4+ T cells (**Supplementary Fig. 7B**). In contrast, even fewer CD8+ T cells (1.3%) or DN T cells (0.88%) expressed FoxP3 (**Supplementary Fig. 7A, C**). Therefore, very few of the tissue-infiltrating CD4+ T cells in lupus nephritis are potentially Tregs.

We additionally identified T follicular helper-like (Tfh) cells based on the combination of PD1 and ICOS expression by CD4+ T cells (*29, 30*). 5.05% of the CD4+ T cell compartment were PD1+ICOS+FoxP3-. Previous investigations have consistently associated PD1 expression with Tfh-like cells (including T peripheral helper cells) but not necessarily ICOS. Therefore, we applied a less stringent definition of PD1+ICOS+/- FoxP3-to identify this cell subset. This Tfh phenotype was associated with roughly 30% of the CD4+ T cells (**Supplementary Fig. 7B**). Although PD1+ICOS-FoxP3-CD4+ T cells could be interpreted as exhausted, we chose to use the more expansive Tfh definition in our subsequent analysis.

### Organization of inflammation across whole biopsies

We next probed potential interacting partners of Tfh, Treg, and Tex cells by identifying the class of their nearest neighbors. Most Tregs are closest to other Tregs and other CD4+ T cells (**Supplementary Fig. 7D**). In contrast to the expectation that Tfh would primarily be in close proximity with CD20+ B cells, Tfh had other CD4+ T cells as their most frequent neighbor, followed by other Tfh (**Supplementary Fig. 7E**). Exhausted CD8+ T cells were most frequently found near other exhausted CD8+ T cells, followed by CD4+ T cells and CD8+ T cells (**Supplementary Fig. 7F**). Overall, these data demonstrate that across biopsies there is a tendency for the clustering of similar cells together.

Cell neighborhoods in the HMP dataset were then defined using DBSCAN with a distance cutoff of 50 pixels, roughly 10 µm. Most neighborhoods detected in the HMP dataset were small (**Fig. 7A**). However, without the constraint of discrete fields of view, we were able to capture larger neighborhoods, with a maximum neighborhood size of 273 cells, relative to the 147 cell maximum in the HR dataset. Given the association of CD4-T cell enriched neighborhoods with ESRD+ patients in the HR data, we investigated similar CD4-neighborhoods in the HMP data. We classified CD4-neighborhoods as those that had 1) less than 20 cells and 2) ≥25% of their cells were either CD8+ or DN T cells, as these criteria captured 99.1% of the CD4-neighborhoods observed in the HR data (**Fig. 7B**). A strong majority of the cells in these neighborhoods are CD4-T cells, including 26% DN T cells and 34.2% CD8+ T cells (**Fig. 7C**). There was a weak negative correlation (R=-0.35) between the number of DN T cells and CD8+ T cells in these neighborhoods, suggesting that DN and CD8+ T cells are not proportionally represented in a given neighborhood.

**Fig. 7.**
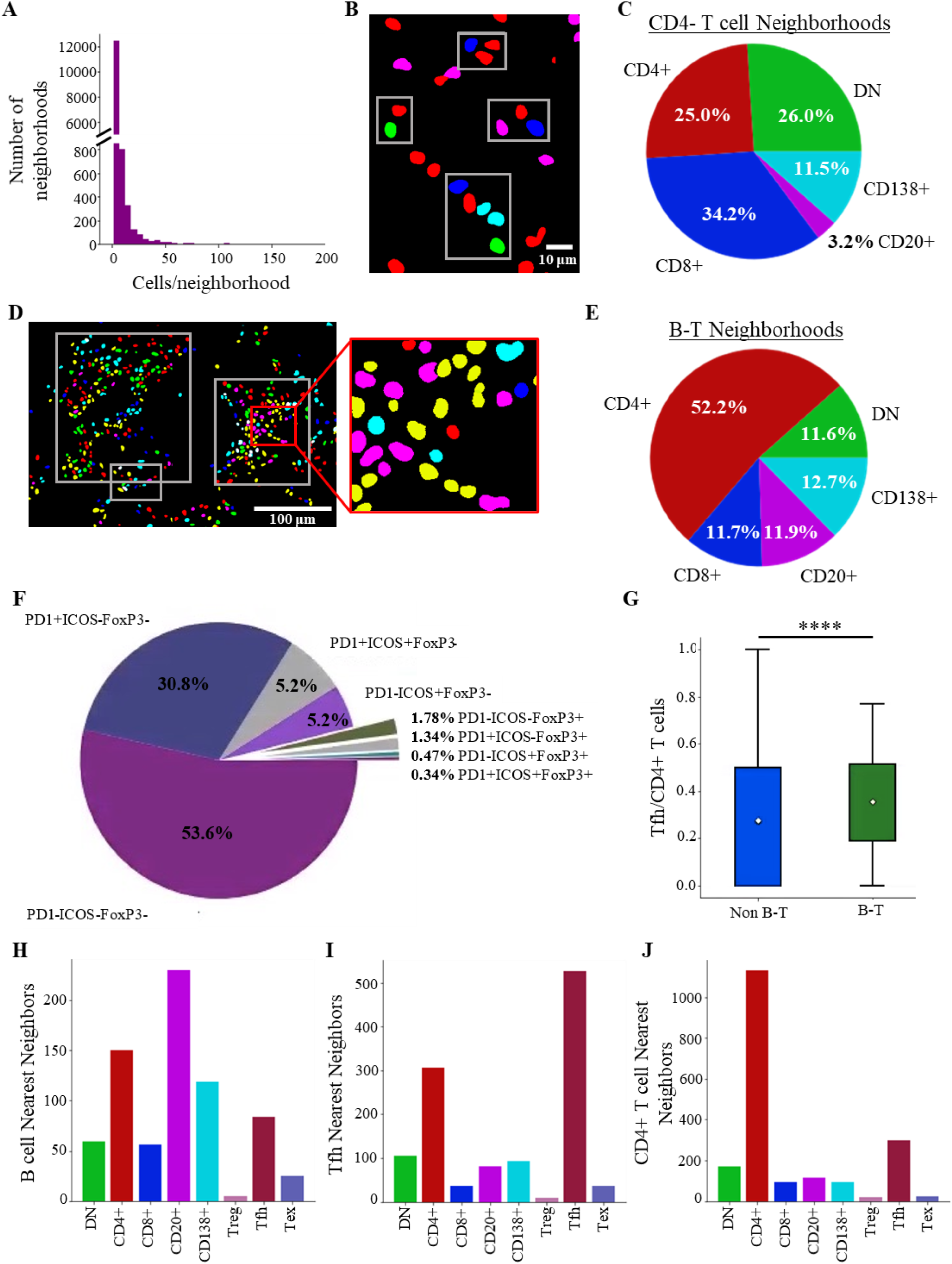
Identification of distinct CD4- and B-T neighborhoods; A) Distribution of sizes of all cell neighborhoods in the HMP dataset. B) Representative CD4-clusters, red=CD4+ T cells, blue=CD8+ T cells, green=DN T cells. C) Distribution of the five main lymphocyte classes in the CD4-T cell neighborhoods. D) Representative B-T aggregates (outlined by white boxes), green=DN, red=non-Tfh CD4+, yellow=Tfh; blue=CD8+, magenta =CD20+, cyan=CD138+ cells. E) Distribution of the five base classes of lymphocytes in B-T neighborhoods. F) Distribution of CD4+ T cell phenotypes in B-T neighborhoods. G) Comparison of proportion of CD4+ T cells that are Tfh in identified B-T aggregates and non B-T aggregates (Mann-Whitney U Test, p =1.9×10^−6^). The nearest neighbors of H) CD20+ B cells, I) Tfh, and J) CD4+ T cells within B-T aggregates.

Large B-T neighborhoods were defined by the set of three criteria (described above) that captured most of the large, lymphocyte-rich neighborhoods in the HR data. Of nearly 14,000 neighborhoods in the HMP dataset, 111 met these criteria (representative clusters in **Fig. 7D)**. Within these B-T neighborhoods, a vast majority of the lymphocytes were T cells, followed by similar proportions of B cells and plasma cells (**Fig. 7E**). Tfh cells made up 36% of CD4+ T cells in B-T neighborhoods (**Fig. 7F)**, a significant enrichment compared to non-B-T neighborhoods (p = 1.9×10^−6^, Mann-Whitney) **(Fig. 7G)**. As observed across whole biopsies, within these B-T neighborhoods, homotypic proximity predominated. B cells were located near other B cells, followed by plasma cells and CD4+ T cells (**Fig. 7H**). Tfh cells in these neighborhoods were most often near other Tfh cells, while overall Tfh were near unspecified CD4+ T cells (**Fig. 7I, Supplementary Fig. 7E**). Unassigned CD4+ T cells in B-T neighborhoods were also most likely to be found near other CD4+ T cells, followed by Tfh cells (**Fig. 7J**). These data suggest that cellular neighborhoods first identified in discrete fields of view, are an organizing principle across whole biopsies.

## DISCUSSION

Canonically, lupus nephritis is thought of as arising from a systemic break in B cell tolerance leading to glomerular antibody deposition and inflammation. This model, supported by large bodies of evidence in both humans and mice (*10, 31*), has led to clinical trials targeting B cells and Tfh cells (*32-38*). However, these efforts have yielded either incremental or no improvement over the standard of care. By quantifying cellular organization within confocal microscopy images using deep learning algorithms, we demonstrate that high regional B cell density is associated with a good prognosis. Rather, it is CD4-T cell populations, including CD8+, γδ, and other DN T cells, that are associated with refractory disease and progression to renal failure.

The CD4-population was surprisingly heterogeneous. As expected, CD8 expressing cells were common. However, over 40% of the CD4-cells did not express CD8. Of these approximately 50% expressed the γδ TCR. Re-examination of the AMP data confirmed the presence of intrarenal γδ T cells within the DN T cells. We also observed a substantial population of CD4-CD8-δ-T cells. These cells appear similar to previously described DN T cells which arise from CD8+ self-reactive T cells that have downregulated CD8 expression (*39, 40*).

It remains to be determined whether a specific CD4-T cell population is associated with progression to renal failure or if these populations share pathogenic roles. Certainly, both CD8 and γδ T cells can be cytolytic and might provide complementary recognition of different classes of autoantigens (*41-44*). Intrarenal γδ T cells have been implicated in chronic renal disease, though the mechanism is unclear (*45, 46*). The function of DN T cells is not known, but they might retain cytolytic activity, as they are derived from CD8 T cells. Alternatively, they could be a source of inflammatory cytokines (*39*). Resolving heterogeneity in these CD4-T cell populations is necessary to identify the populations most closely linked with ESRD.

In addition to yielding information on cell frequencies in tissue, our analytic pipelines provided precise positions of all cells assayed in the biopsy. This allowed us to define cellular neighborhoods and extract quantitative features including neighborhood size, shape and cell constituency. Unsupervised clustering revealed that in individual patients, small neighborhoods of CD4-T cells were associated with progression to ESRD. These data suggest that understanding immune cell architectures, even in relatively small patient sample cohorts, can identify prognostically important mechanisms.

Our data identify CD4-T cells, including CD8+, γδ, and DN T cells as potentially important therapeutic targets. This association was particularly striking in patients that presented in renal failure. It is possible that the inflammatory phenotype observed in these patients was not a primary state but arose as a secondary consequence of renal damage and scarring. However, two of these patients (patients 51 and 55) had high densities of CD4-T cells, high activity indices, and relatively low chronicity scores, suggesting that infiltrating CD4-T cells can precede substantial renal damage. These data suggest that patients exhibit distinct, prognostically meaningful, intra-renal inflammatory trajectories.

Unfortunately, in contrast to the B cell:Tfh axis, we have limited therapeutic options that specifically target these T cell populations. One of the few classes available are calcineurin inhibitors (CNIs). The recent success of the CNI voclosporin in treating some patients with lupus nephritis is promising (*47*). We propose that stratifying patients by the constituency and organization of their renal inflammation might identify those most likely to benefit from the addition of T cell targeting therapies such as voclosporin.

Dense regions of B cells were surprisingly associated with a good renal outcome. It is possible that B cells and subsequent local antibody secretion are somewhat benign compared to other pathological processes. Alternatively, conventional therapies might be relatively effective against B cells and plasma cells. Indeed, most of our patients were treated with high-dose steroids and induced with cytotoxic therapies, most often mycophenolate. These therapies have been demonstrated to deplete B cells and plasma cells (*48, 49*).

Large neighborhoods of cells were enriched in both B cells and CD4 T cells, including putative Tfh cells. These neighborhoods have similar features to the T:B aggregates described previously (*15*). However, our HMP image analysis indicated that these structures are more complex, containing other cell types including other CD4+ T cell and PC populations. These findings cohere with previous studies that suggest an underlying architecture to these large neighborhoods (*13*). Further work will be needed to understand the rules by which these different neighborhoods organize with respect to each other, and the underlying biological processes governing their organization.

Interestingly, pDC prevalence was not associated with renal outcome in our HR dataset. As sources of IFNα, they have been postulated to play a central role in disease pathogenesis (*50-53*). However, the outcomes of clinical trials of anti-IFNα antibodies in lupus have been modest (*54, 55*). In contrast to pDCs, mDCs were depleted in those patients that presented in renal failure. Therefore, they might have a role in organ tolerance even in the context of severe inflammation. It should be noted that the interstitial mDCs we quantified appeared different than the periglomerular inflammatory mDCs that have been recently described in lupus nephritis (*56*). Furthermore, CD11c is a marker that can capture a range of cells in addition to mDCs, including macrophages and even age-associated or double negative B cells (*57, 58*). In our CNN hierarchy, putative cells expressing both CD20 and CD11c would be classified as B cells; such occurrences were rare. Further work, with multiple additional cell markers, will be needed to resolve the complexity of the CD11c populations.

Using deep learning and other artificial intelligence algorithms, we achieved robust and accurate cell detection across multiple lupus nephritis image datasets. This enabled a detailed, accurate spatial analysis of *in situ* immunity in lupus nephritis samples. Several computer vision methods were implemented to establish an analytical pipeline that addressed experimental, biological, and technical limitations. CNNs trained for instance segmentation detected and classified several immune cell classes with high fidelity not only in sparsely populated images, but also in densely packed images. We also trained and implemented CNNs for rapid and robust image and object filtering to optimize immune cell calling in full-biopsy sections. This included a network trained to discriminate between image tiles containing positive cell signal and tissue autofluorescence. We also trained a network to segment tubules in order to reject false-positive plasma cell predictions. By combining these CNNs with thresholding and image registration techniques, we automatically mapped several immune cell classes to full-biopsy sections enabling a robust spatial analysis of *in situ* autoimmunity.

We implemented artificial intelligence to quantify immune cell populations in human tissue, thereby extracting rich, non-biased, spatio-cellular data that allowed identification of unexpected pathogenic mechanisms. Even in a relatively small longitudinal cohort, we were able to resolve patient heterogeneity to identify putative pathogenic processes. Remarkably, specific cell densities provided powerful insights into disease pathogenesis. Understanding how these populations were organized into neighborhoods provided predictive power in individual patients. Furthermore, this work revealed fundamental insights to the structural organization of inflammation that began to identify organizing principles. Further studies of the complexity, heterogeneity, and organization of *in situ* inflammation will yield a more quantitative understanding of human autoimmunity. Such knowledge is critical for interpreting and applying the wealth of knowledge we have gained from animal models. It is also likely to identify both new therapeutic targets and those patients in which specific strategies are likely to be beneficial.

## MATERIALS AND METHODS

### Sample staining and image acquisition: High-resolution dataset

Formalin-fixed paraffin-embedded (FFPE) kidney biopsies from 55 lupus nephritis patients with at least two years of clinical follow-up were obtained from the University of Chicago Human Tissue Resource Center. FFPE sections were de-paraffinized and treated with a citric acid buffer (pH 6.0) for antigen retrieval and blocked with serum prior to antibody staining. Samples were stained with indicated specific antibodies (**Supplementary Table 2**) and imaged on a Leica SP8 laser scanning confocal microscope at 63x magnification. Image ROIs were collected in tissue regions with identifiable CD3 signal. Collected images were 1024 × 1024 pixels x 6 channels with a 0.1058 µm pixel size.

### Staining and image acquisition: Highly multiplexed dataset

Samples were stained using a strip and reprobe procedure in which 5 µm thick sections of FFPE biopsy sections were iteratively stained according to a procedure outlined by (*26*). Sections were deparaffinized and stained with a combination of primary antibodies and secondary antibodies conjugated with AlexaFluor 488, 546, and 647 fluorophores (**Supplementary Table 2**). DAPI was included in every iteration of staining. Each round of staining, samples were imaged using a Caliber ID RS-G4 large-format confocal microscope at a magnification of 63x, resulting in a pixel size of 221 nm. After each round of imaging, samples were stripped as described (*26*) then re-probed with a new set of primary and secondary antibodies and re-imaged until the full marker panel had been imaged.

### Staining and image acquisition: γδ T cells

Eight lupus nephritis kidney biopsies were stained for CD3, CD4, CD8, TCRδ, and DAPI. Inflamed regions were imaged on the Leica Stellaris 8 confocal microscope, with 40x magnification and a pixel size of 0.225 um. 281 ROIs (35±19 per sample) were obtained, and then post processed with background subtraction, de-speckling and contrast adjustment using ImageJ. Cells in these images were quantified by manual count.

### Automatic cell detection and segmentation: High resolution dataset

Two instances of a Mask R-CNN architecture (*59*) were trained to detect and segment cells in this dataset. One instance was trained to detect three classes of lymphocytes (B cells, CD3+CD4-T cells, and CD3+CD4+ T cells). A second instance was trained to detect two classes of DCs (pDCs and mDCs). For this work, ResNet-101 was used as a backbone for the FPN. Networks were trained with a learning rate of 0.01. Training, validation, and testing data were generated by a single expert. All manually segmented images from a given patient were relegated to either the training set (273 images, 80%), validation set (34 images, 10%), or test set (35 images, 10%). Training progress was monitored using Tensorboard and training was stopped after cell recall stopped improving for all classes. Precision (Eq.1), recall (Eq. 2), and F1-score (Eq. 3) were used to evaluate network performance. All computation for the HR dataset was performed using resources at the University of Chicago Research Computing Center. Each instance of Mask R-CNN was trained on a single GPU compute node containing four Nvidia GPUs with 12GB memory per card, 28 Intel E5-2680v4 CPUs at 2.4 GHz, and 64 GB of system memory. A batch size of 4 images was used for training, distributed across the four GPU cards. Networks were trained to the point at which the recall for all cell classes stopped increasing.

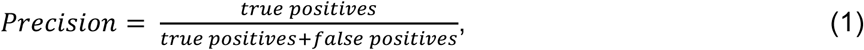

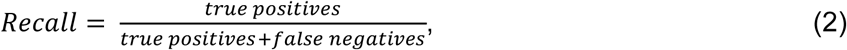

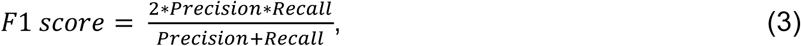

### Automatic cell detection and segmentation: Highly multiplexed dataset

Image strips (1024 x N pixels) from Caliber ID microscope were stitched together using cross-correlation of image patches at the strip boundaries. These single-channel composites were then aligned with the DAPI channel from the first round of imaging, again through cross-correlation of image patches. Multi-channel composites were then broken into 512 × 512-pixel image tiles. All DAPI tiles were passed through a simple image intensity filter to determine if tissue was present at a given location. All tiles at a given location were filtered out of proceeding analyses if the DAPI filter revealed no tissue at that location. This resulted in over unique 18,000 tile locations, with each location containing nine unique stains.

All image tiles that contained tissue were further filtered for the presence of lymphocyte signal (CD3 or CD20) using a custom 18-layer CNN trained to classify images based on positive cell signal. Instance segmentation of cells was split into two tasks: instance segmentation of T cells (also referred to as dual-marker detection) and instance segmentation of B cells (also referred to as single-marker detection). Instance segmentation performance was evaluated using precision (Eq. 1), recall (Eq. 2), and F1-score (Eq. 3).

#### T cell segmentation

An instance of Mask R-CNN was pre-trained to segment single positive (CD3+ CD4-) and double positive (CD3+CD4+) T cells in 512 × 512-pixel image patches from T cell image stacks (CD3/CD4/DAPI) from the HR dataset. This pre-trained network was fine-tuned with a small set of 211 T cell image stacks from the highly multiplexed dataset. The highly multiplexed fine-tuning image sets contained CD3/CD4/DAPI, CD3/CD8/DAPI, and CD3/ICOS/DAPI image stacks. This fine-tuning set was split into training (169 images, 80%), validation (21 images, 10%), and testing (21 images, 10%) sets, with all images from a given patient confined to a specific set. For fine-tuning, weights were permitted to adjust for all convolutional, max-pooling, and fully connected layers of the pre-trained Mask R-CNN. The fine-tuned T cell network was used to make predictions for CD3/CD4/DAPI, CD3/CD8/DAPI, CD3/ICOS/DAPI, and CD3/PD1/DAPI images. The trained dual-marker network had an average F1-score of 0.85 on all single-positive cell predictions (i.e., CD3+CD4- and CD3+CD8-) and an average F1-score of 0.92 on all double-positive cell predictions (i.e., CD3+CD4+ and CD3+CD8+).

#### B cell segmentation

An instance of Mask R-CNN was pre-trained to segment B cells in 512 × 512-pixel image patched generated from B cell image stacks (CD20/DAPI) from the HR dataset. This pre-trained network was fine-tuned with a set of 79 B cell image stacks from the HMP dataset. This fine-tuning set was split into training (63 images, 80%), validation (8 images, 10%), and testing (8 images, 10%) sets, with all images from a given patient confined to a specific set. For fine-tuning, the weights were permitted to be adjusted for all convolutional, max-pooling, and fully connected layers of the pre-trained Mask R-CNN. The fine-tuned B cell network was used to make predictions for B cell (CD20) and plasma cell images (CD138). The trained single-marker network had and average F1-score of 0.87.

### Tubule segmentation

An instance of Mask R-CNN was trained to segment tubular structures, including blood vessels and tubules, in the HMP dataset. 300 DAPI tiles (512 × 512 pixels) from 18 patients were manually annotated by a single expert. Manually segmented images were separated into training, validation, and testing sets as follows: 240 images in the training set (80%), 30 images in the validation set (10%), and 30 images in the test set (10%). Data augmentation consisted of random horizontal and vertical flips and rotations. Performance of the tubule segmentation network was assessed at the pixel level, with the trained network yielding an average recall (Eq. 2) of 0.74 and an average precision (Eq. 1) of 0.79 on the test set of tubule images.

All computation associated with the HMP dataset was performed on the MEL computational server in the Radiomics and Machine Learning Facility at the University of Chicago. MEL contains 256 Xeon Gold 6130 CPU cores, 3TB of DDR4 ECC RAM memory, 24TB of NVMe SSD storage, and 16 nVidia Tesla V100 32GB GPU accelerators.

### Defining cell neighborhoods through density-based clustering

Cells in both datasets were assigned to clusters using the sklearn (version 0.23.2) implementation of Density Based Spatial Clustering of Applications with Noise (DBSCAN) (*25*), using an epsilon of roughly 10 µm, corresponding to 100 pixels in the HR dataset and 50 pixels in the HMP dataset and a minimum cluster size of two. In the HR dataset, twenty-four features of cellular constituency and cell/neighborhood shape, were extracted for each cluster, and K-means clustering was then applied to define classes of neighborhoods. The neighborhoods were split into six classes as determined ideal by bootstrapping cluster descriptors including the within cluster sum of squares (WCSS) and the delta WCSS. The types were characterized using a leave-one-out t test to identify which features of each type of neighborhood distinguished it from the other neighborhoods. In this procedure, the current cluster of reference was treated as the alternative group, all remaining clusters were then binned together as the reference group, then a t-test for was applied for all features used to describe the neighborhoods.

### Spatial Analyses

All other spatial analyses were performed using Python (3.7.9) and the following packages: pandas (1.2.2), numpy (1.19.2), sklearn (0.23.2), scipy (1.6.1), and tifffile (2021.1.14). Plotting was performed with matplotlib (3.3.2) and seaborn (0.11.1). The nearest neighbors calculation was performed by iterating through every cell in the dataset and identifying the class of the closest cell by centroid-to-centroid distance.

In the HMP dataset, coordinates of the cells in the tiles were adjusted to a composite-level coordinate system by shifting the tile-level coordinates based on the location of the tile in the composite. All subsequent calculations around the distribution of cells in tissue were based on these composite-level locations.

### RNA sequencing analysis

Single-cell RNA-seq data for human lupus nephritis tissue were obtained from the ImmPort repository (accession code SDY997, “SDY997_EXP15176_celseq_matrix_ ru10_molecules.tsv” raw data file). Quality control was performed according to the original paper (*16*), such that cells were removed from the analysis if they expressed <1,000 or >5,000 genes, or if more than 25% of the total unique molecular identifiers (UMI) mapped to mitochondrial genes. Gene expression values were normalized to library size (UMI count per million) and scaled by log2. Clustering implemented in Seurat 3.2.2 and canonical marker expression were used to identify cellular subsets. T cells were analyzed if they were assigned to the “Naïve T” or “CTL” clusters. T cells were categorized based on *CD4, CD8A*, and *CD8B* expression. Cells were categorized as “CD4” when they had detectable expression of *CD4* transcripts but no *CD8A* or *CD8B*. They were instead categorized as “CD8”, when they had detectable *CD8A* and/or *CD8B* with no *CD4* transcripts. Cells were categorized as DP (double-positive) or DN (double-negative) when they had both/neither *CD4* and/nor *CD8A/B*. t-SNE was performed by Rtsne (0.15). Plots were generated by ggplot2 (3.3.2) and ggridges (0.5.2).

## Acknowledgments

Special thanks to Karen Drukker, PhD for her input on statistical analyses, Chun-Wai Chan, MS for computational support, and the University of Chicago Research Computing Center for additional computational support and resources. All imaging was performed at the University of Chicago Integrated Light Microscopy Facility.

## Author Contributions

Conceptualization: RA, MD, MRC

Methodology: RA, MD, JA, MV, GC, YA, MG, MRC

Software: RA, MD, GC, YA

Investigation: RA, MD, JA, MV, GC, YA

Resources: AC, KK, EP, CO

Funding acquisition: MRC, MG

Supervision: MRC, MG

Writing – original draft: RA, MD, JA, MV, GC, YA, MRC

Writing – review & editing: RA, MD, JA, MV, GC, YA, MG, MRC

## Data and materials availability

Code used for the cellular segmentation and spatial analysis can be found at the following repository: https://github.com/durkeems13/LN_image_analysis_STM.git Raw image data is available upon request to the corresponding author.

