## Supplementary Materials for "In lupus nephritis, specific *in situ* inflammatory states are associated with refractory disease and progression to renal failure"

Supplementary Figures and Tables

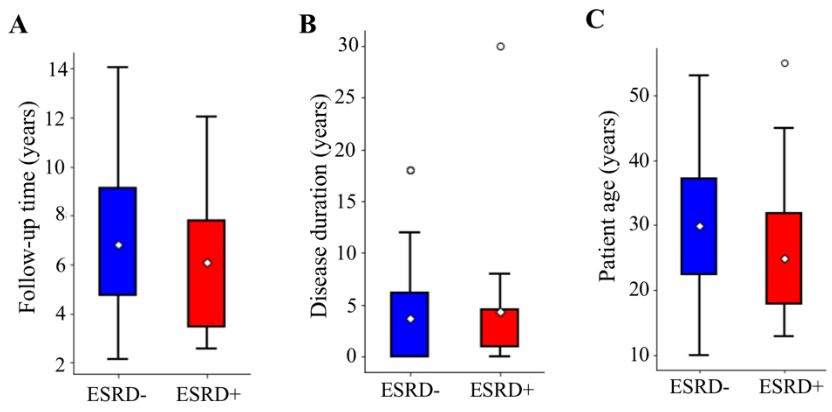


**Supplementary Figure 1. Clinical characteristics of patient cohort.** ESRD+ and ESRD- patients did not differ in A) Duration of follow up period (p=0.678), B) Disease duration (p=0.819), or C) Patient age (p=0.096). (Mann-Whitney U Test with Bonferroni correction for multiple comparisons).

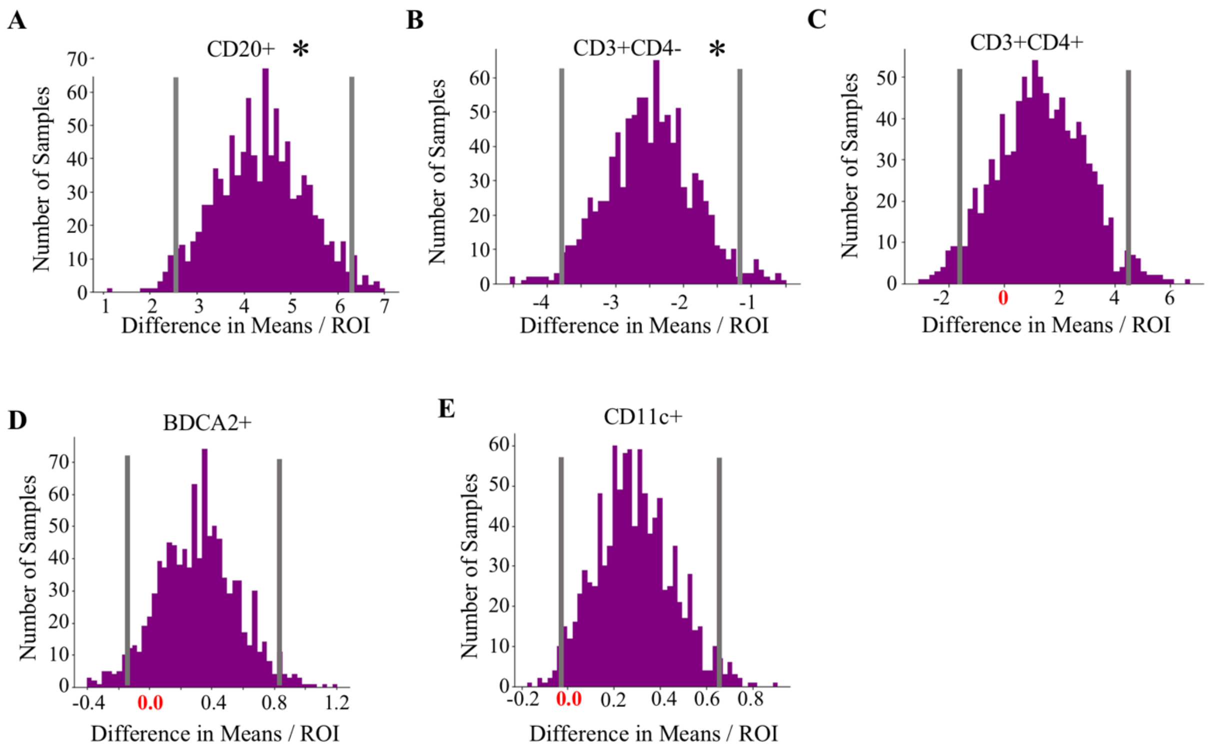


**Supplementary Figure 2**. **Distribution of the difference in means for indicated cells/ROI comparing ESRD+ vs ESRD-.** Difference in means for all iterations of bootstrapping are shown for A) CD20+, B) CD3+CD4-, C) CD3+CD4+. D) CD11c+, E) BDCA2+ cells. Vertical grey lines denote 95% confidence intervals, stars denote the confidence intervals that do not overlap with zero.

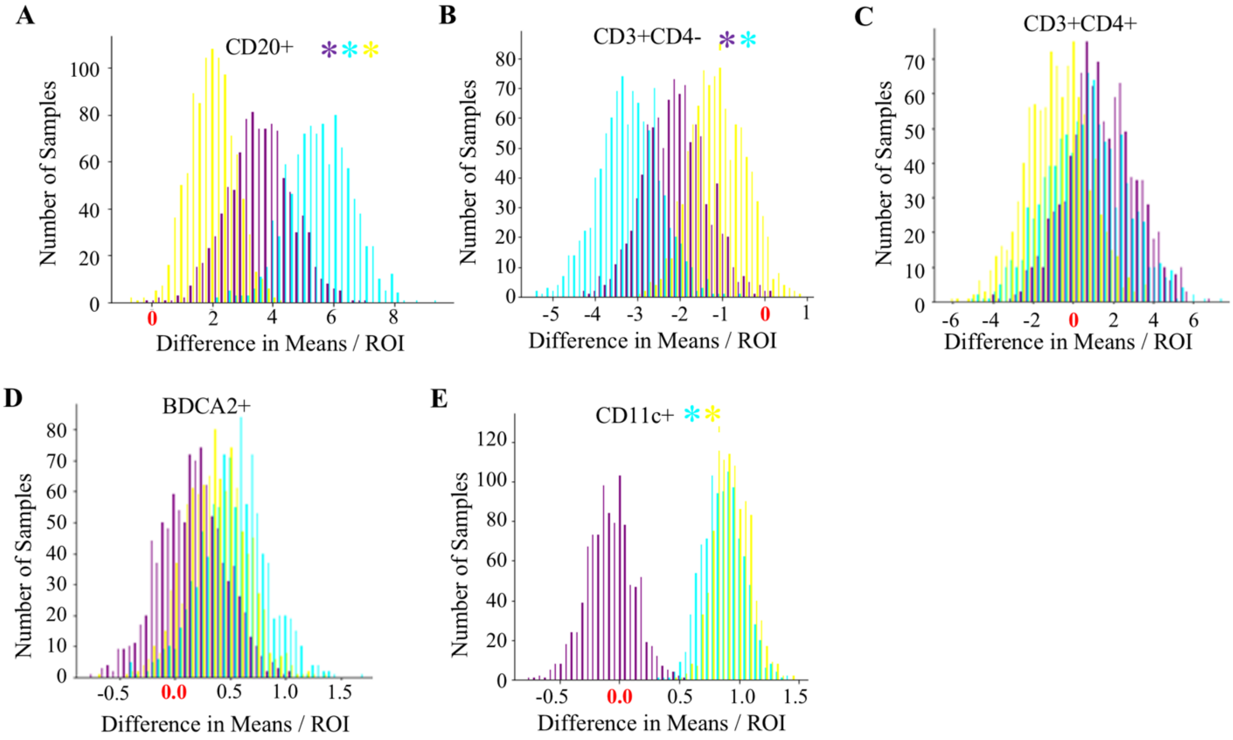


**Supplementary Figure 3. Distribution of the difference in means for indicated cells.** Difference in means for each bootstrapping iteration comparing ESRD+ vs ESRD- (purple), ESRD+ vs ESRD current (cyan), and ESRD- vs current (yellow) for A) CD20+, B) CD3+CD4-, C) CD3+CD4+. D) CD11c+, E) BDCA2+ cells. Stars denote the confidence intervals that do not overlap with zero, colors correspond with the comparison.

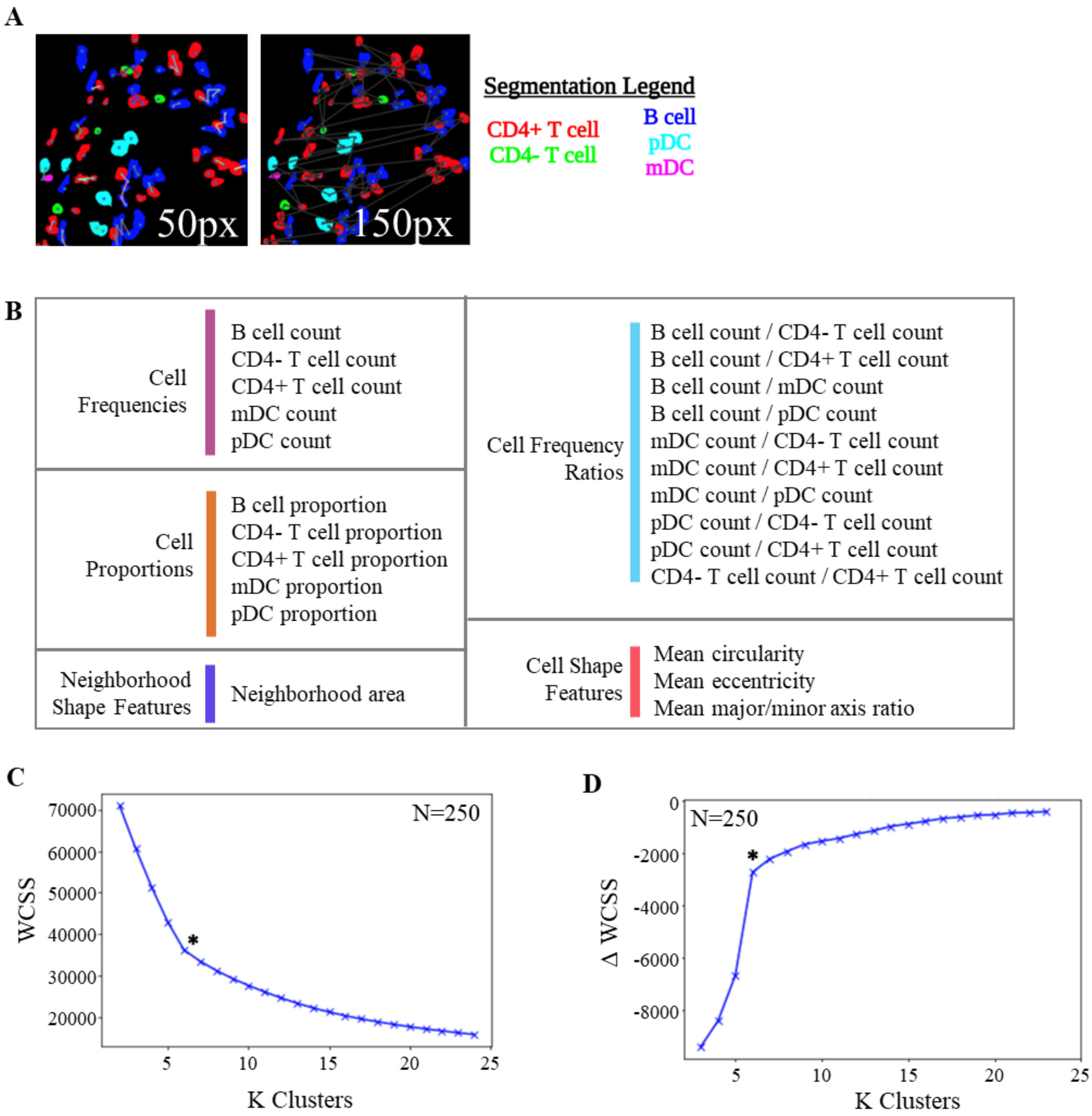


**Supplementary Figure 4. Definition of cell neighborhoods in the HR dataset.** A) Representative outputs of DBSCAN algorithm with varying distance cutoffs (50, 100, and 150 pixels), B) 24 features used to define types of aggregates, results of bootstrapping method for determination of optimal cluster number using C) within cluster sum of distances squared (WCSS) and D) delta WCSS. The optimal cluster hyperparameter used for downstream analyses (k = 6), is denoted by an asterisk.

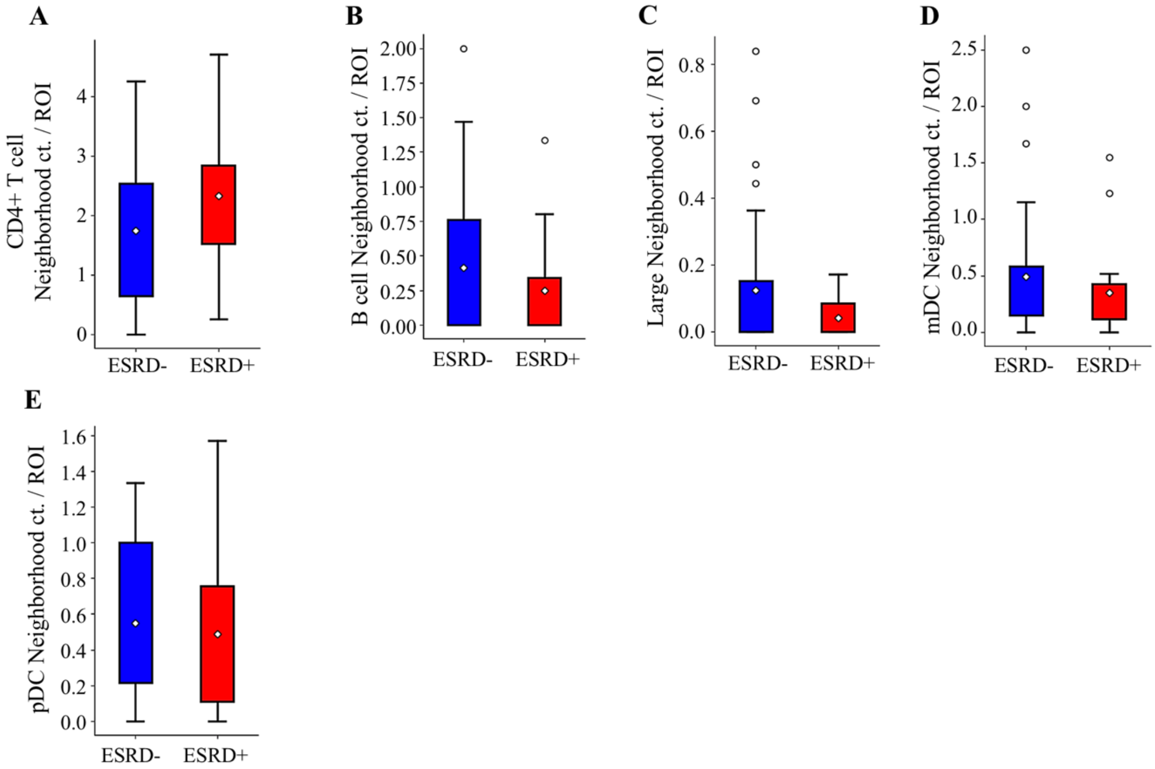


**Supplementary Figure 5. Other cellular neighborhoods not associated with renal failure.** The abundance of neighborhoods between the patient cohorts, normalized by the number of ROIs per patient, is compared by Mann-Whitney U Test, with a Bonferroni correction for A) CD4+ T cell neighborhoods, B) B cell neighborhoods, C) Large neighborhoods, D) mDC neighborhoods, E) pDC neighborhoods

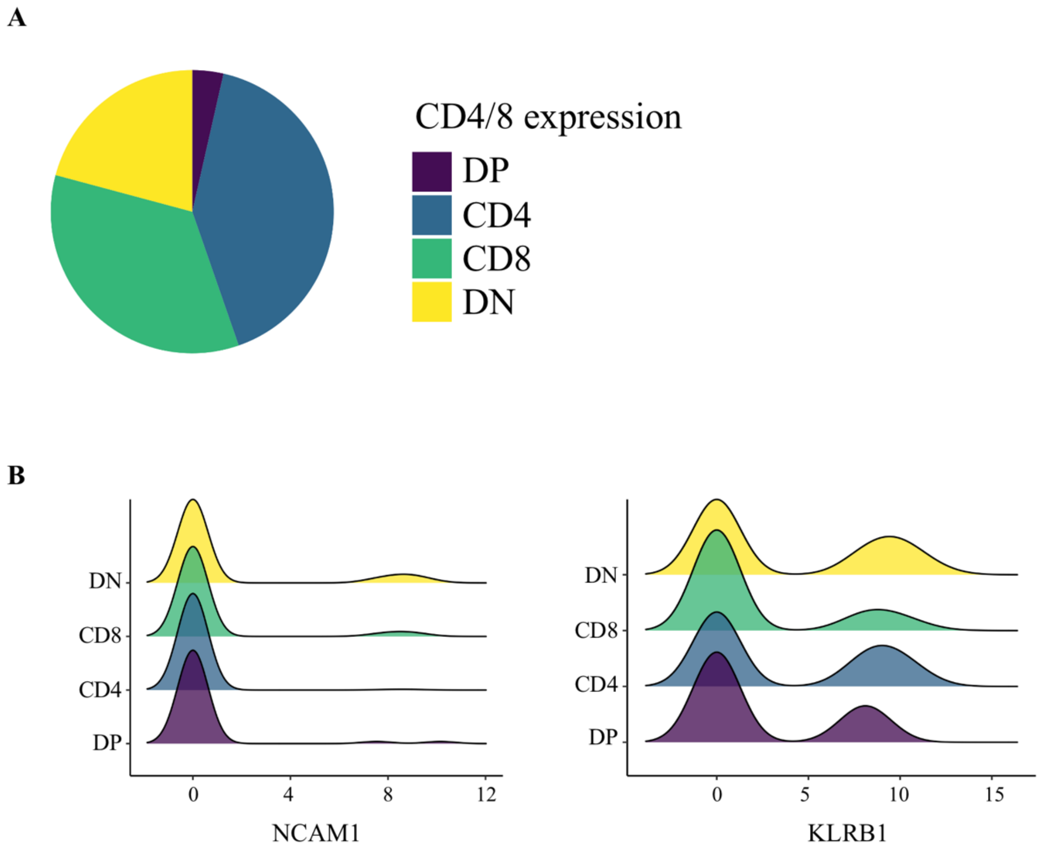


**Supplementary Figure 6. DN T cells are found in scRNA-Seq LN data.** A) Distribution of *CD4/8A/8B* expression in T cell population. DP: double-positive, DN: double-negative; B) Density plots showing distribution of *NCAM1* and *KLRB1* expression.

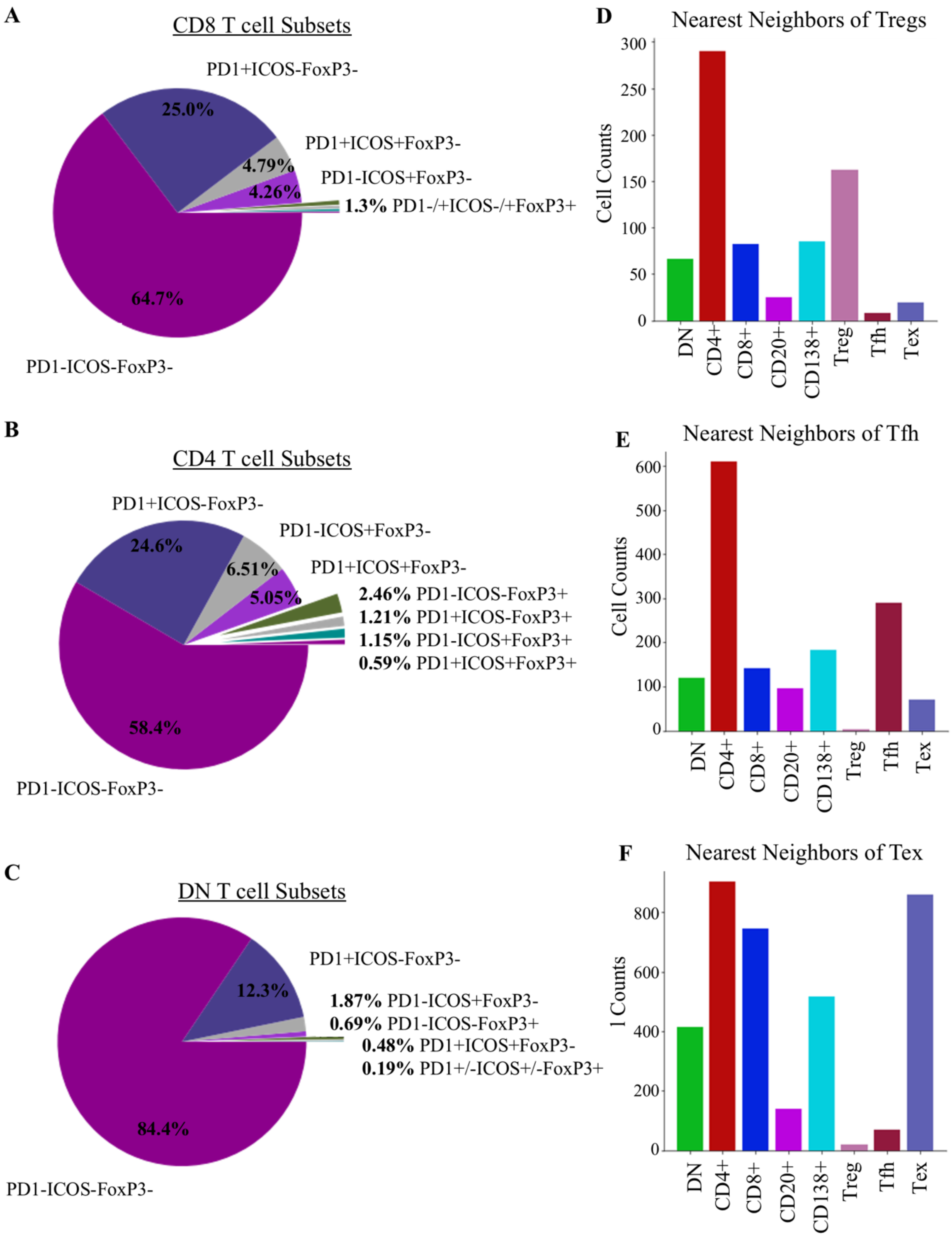
A
CD8 T cell Subsets
PD1+ICOS-FoxP3-
25.0%
PD1+ICOS+FoxP3-
4.79%
PD1-ICOS+FoxP3-
4.26%
1.3% PD1-/+ICOS-/+FoxP3+
64.7%
PD1-ICOS-FoxP3-
B
CD4 T cell Subsets
PD1+ICOS-FoxP3-
24.6%
PD1-ICOS+FoxP3-
6.51%
PD1+ICOS+FoxP3-
5.05%
2.46% PD1-ICOS-FoxP3+
1.21% PD1+ICOS-FoxP3+
1.15% PD1-ICOS+FoxP3+
0.59% PD1+ICOS+FoxP3+
58.4%
PD1-ICOS-FoxP3-
C
DN T cell Subsets
PD1+ICOS-FoxP3-
12.3%
1.87% PD1-ICOS+FoxP3-
0.69% PD1-ICOS-FoxP3+
0.48% PD1+ICOS+FoxP3-
0.19% PD1+/-ICOS+/-FoxP3+
84.4%
PD1-ICOS-FoxP3-
D
Nearest Neighbors of Tregs
Cell Counts
0
50
100
150
200
250
300
DN
CD4+
CD8+
CD20+
CD138+
Treg
Tfh
Tex
E
Nearest Neighbors of Tfh
Cell Counts
0
100
200
300
400
500
600
DN
CD4+
CD8+
CD20+
CD138+
Treg
Tfh
Tex
F
Nearest Neighbors of Tex
l Counts
0
200
400
600
800
DN
CD4+
CD8+
CD20+
CD138+
Treg
Tfh
Tex

**Supplementary Figure 7. CD8, CD4, and DN T cell phenotypes in LN.** Distribution of secondary marker (ICOS, PD1, FoxP3) expression in A) CD8+, B) CD4+ and C) DN T cells, Distribution of nearest neighbors of D) Treg (CD3+CD4+PD1-ICOS-FoxP3+) cells, E) Tfh (CD3+CD4+PD1+ICOS+/-FoxP3+) cells, and F) Tex (CD3+CD4+PD1+ICOS-FoxP3-).

**Supplemental Table 1. Select clinical feature from patients in the HR dataset.** (Table attached as an excel sheet)

**Supplemental Table 2. Reagents for sample staining.** Anti-CD20(Agilent, M0755), anti-CD4(Abcam, ab133616), anti-CD3(BIO-RAD, MCA1477), anti-BDCA2(R&D, AF1376), and anti-CD11c (Abcam, ab52632) were applied for high-resolution images. For high-dimensional images, more antibodies were added, including anti-CD8(Abcam, ab17147), anti-ICOS(Abcam, ab105227), anti-PD1(Abcam, ab52587), anti-Foxp3(Invitrogen, 14-4776-82), anti-CD138(Invitrogen, MA5-12400), anti-MX1(R&D, AF7946), and anti-TCRδ(Santa Cruz, sc-100289). DAPI (Invitrogen, H3570) was used for nuclear staining.

| **Target** | **Vendor** | **Catalog#** |
|---|---|---|
| CD20 | Agilent | M0755 |
| CD4 | Abcam | ab133616 |
| CD3 | BIO-RAD | MCA1477 |
| BDCA2 | R&D | AF1376 |
| CD11c | Abcam | ab52632 |
| CD8 | Abcam | ab17147 |
| ICOS | Abcam | ab105227 |
| PD1 | Abcam | ab52587 |
| FoxP3 | Invitrogen | 14-4776-82 |
| CD138 | Invitrogen | MA5-12400 |
| MX1 | R&D | AF7946 |
| TCRδ | Santa Cruz | sc-100289 |
| DAPI | Invitrogen | H3570 |
